# Miniaturized Traceless Staudinger Ligation Enables Direct-to-Biology Screening of Potent BRD4 PROTACs

**DOI:** 10.64898/2026.11.07.765033

**Authors:** Marko Mitrović, Leann Schüffner, Saran Aswathaman Sivashanmugam, Stefan Knapp, Kerstin Hiesinger

**Author notes:** These authors contributed equally (Marko Mitrović, Leann Schüffner).

## Abstract

Direct-to-biology (D2B) approaches accelerate PROTAC discovery by enabling biological screening of crude reaction mixtures without intermediate purification. However, the range of chemical transformations compatible with these workflows remains limited. Herein, we establish a miniaturized, base-free traceless Staudinger ligation platform diversifying linker chemistry for a preexisting library of E3-ligand-linker adducts bearing azide moieties. As a proof-of-concept target, BRD4 was chosen, yielding amide-linked BRD4-targeting PROTACs. By combining phosphine-functionalized JQ1 derivatives with 72 selected azide building blocks, reactions were performed on a 5 µL scale in 384-well plates, achieving a mean LC–MS conversion of approximately 65%. Spiking experiments confirmed that the phosphine starting materials had no significant impact on BRD4 degradation, supporting the direct biological evaluation of crude reaction mixtures in the presence of residual starting material. Cellular screening identified 23 hits at 250 nM screening concentration, reducing BRD4 levels to ≤40%. Two resynthesized and purified PROTACs were subsequently validated, displaying pDC_50_ values of 8.2 and 7.0 and a D_max_ of 90% and 96%, respectively. This work establishes traceless Staudinger ligation as a complementary D2B strategy for the rapid discovery of amide-linked PROTACs.

---

Direct-to-biology (D2B) approaches have emerged as valuable tools to expedite hit discovery in medicinal chemistry by enabling the biological evaluation of crude reaction mixtures in in vitro and cellular assay systems (Scheme 1).^1^ This approach is particularly attractive for the development of proteolysis-targeting chimeras (PROTACs), which require a significant synthetic effort due to the considerable number of combinantions of protein of interest ligands (POIs), linkers and linker attachment point variations and E3 ligase and ligand choices.^2^ Owing to their modular architecture, PROTAC optimization requires structural diversification through systematic variation of linker moieties, attachment points and ligands. Given the current lack of rational design strategies and the resulting labor- and resource-intensive efforts, establishing miniaturized synthesis coupled with high-throughput biological evaluation substantially streamlines degrader discovery. A central requirement for D2B synthesis is the availability of robust transformations that proceed efficiently under conditions compatible with subsequent biological evaluation. Amide coupling has been successfully established for miniaturized D2B synthesis and purification-free screening of PROTAC libraries, including nanomole-scale synthesis and high-throughput cellular degradation assays.^3–6^ Nevertheless, expanding the repertoire of D2B-compatible transformations remains desirable to access complementary chemical space for a predefined library of diverse E3 ligands and linkers.

Copper-catalyzed azide–alkyne cycloaddition (CuAAC) represents an exceptionally reliable reaction for D2B applications, given its high chemoselectivity and robustness in parallel synthesis. We previously established a CuAAC-based D2B platform^7^ for PROTAC discovery using 48 azide building blocks from our in-house library^8^. Since then, our azide collection has been further expanded, providing a versatile resource for exploring additional PROTAC chemotypes. However, CuAAC inherently introduces a triazole linkage, motivating the development of complementary transformations that enable the same building blocks to access alternative connecting functionalities.

Traceless Staudinger ligation (TSL) offers an attractive complementary approach by enabling the chemoselective formation of amide bonds from azides and appropriately functionalized phosphines without requiring a metal catalyst (Scheme 1).^9,10^ The reaction proceeds through an iminophosphorane intermediate, followed by intramolecular acyl transfer and hydrolysis to afford the corresponding amide. Notably, Bemis and co-workers previously demonstrated the utility of TSL for the parallel synthesis of PROTAC linker variants.^11^ However, the resulting compounds required purification prior to biological evaluation. Furthermore, comparatively slow reaction kinetics, substrate-dependent conversion, and the formation of phosphorus-containing byproducts pose challenges for implementing TSL in purification-free D2B workflows.

**Scheme 1::**
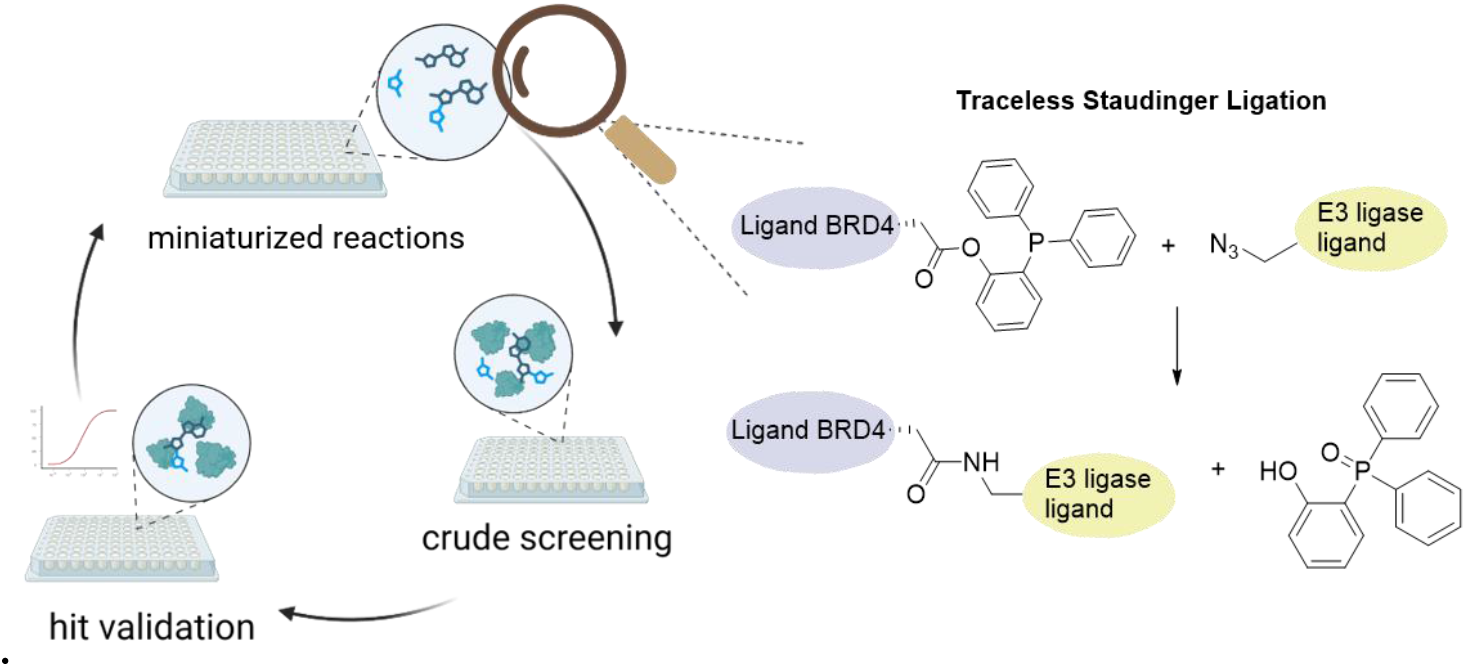
Schematic overview of the miniaturized traceless Staudinger ligation direct-to-biology (D2B) workflow for BRD4 PROTAC discovery.

Herein, we report the development of a miniaturized, base-free TSL platform for the direct biological evaluation of amide-linked Bromodomain-containing protein 4 (BRD4)-targeting PROTACs as proof of concept utilizing TSL in a D2B setting. By combining three phosphine-functionalized derivatives, obtained from the highly potent BRD4 inhibitor JQ1^12^, with 72 selected azides from our in-house library, we generated two series of PROTAC candidates for direct screening of crude reaction mixtures. With a hit-selection threshold of ≤40% remaining BRD4, 22 hits were identified, demonstrating the applicability of TSL as a complementary D2B transformation for PROTAC discovery.

## Results and Discussion

To establish TSL as a complementary transformation for direct-to-biology PROTAC discovery, we designed phosphine-functionalized derivatives of the BRD4-binding ligand JQ1 (Figure 1). The ligation reagents were accessible from commercially available JQ1 carboxylic acid and phosphine-containing alcohol building blocks through short synthetic sequences comprising one step, or two steps when an additional linker was introduced. This modular strategy enabled variation of the connectivity between the BRD4-binding moiety and the reactive phosphine functionality without requiring the development of specialized phosphine reagents. Three phosphine-functionalized JQ1 derivatives, **73, 74**, and **75**, were investigated during reaction development (Figure 1). **75** incorporates a longer linker on the JQ1 side than **74. 73** and **74** differ solely in the connectivity of the phosphine moiety, featuring either a phenyl or benzyl group. Initial optimization was performed using equimolar amounts of phosphine-containing coupling partner **73** and azide **18** in 5 µL reactions conducted in 384-well PCR plates at 40 °C for 18 h. Beyond achieving sufficient chemical conversion, establishing a robust D2B workflow required reaction conditions compatible with accurate microscale liquid handling and subsequent biological evaluation. A particular challenge was the limited solubility of the phosphine-functionalized JQ1 derivatives in THF and water. This limitation was especially relevant at the miniaturized scale, where reagent aliquots of 2 µL were dispensed into individual wells. Pipetting heterogeneous suspensions at such small volumes can compromise dosing accuracy and reproducibility, potentially introducing variation in reagent stoichiometry between reactions. Accordingly, solvent selection was guided not only by ligation efficiency but also by the need to maintain homogeneous reagent solutions. Although more soluble phosphine derivatives have been reported ^13^, we deliberately employed commercially available phosphine-containing alcohol building blocks to preserve the synthetic accessibility and practical transferability of the platform.

**Figure 1.**
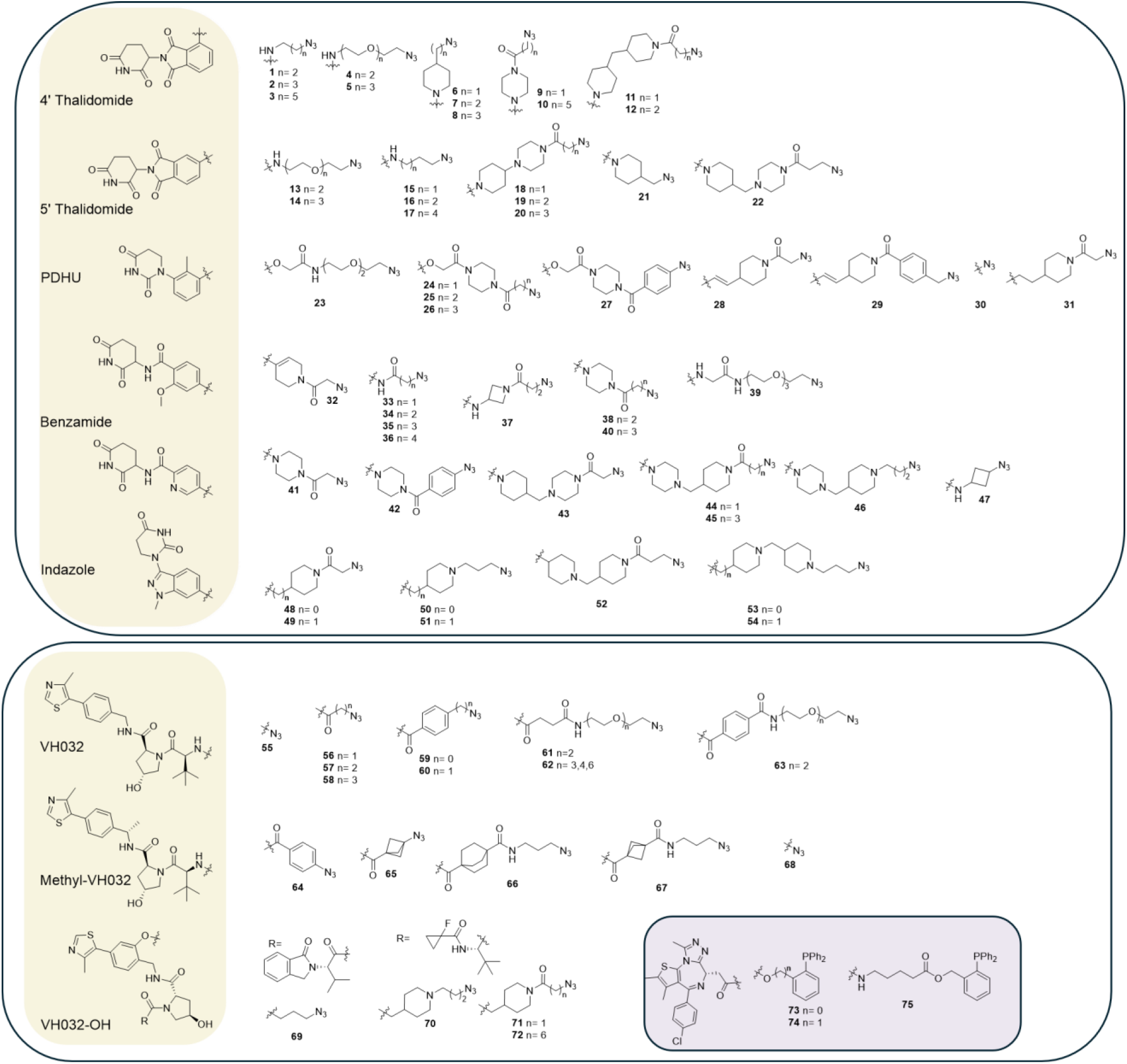
Overview of the azide building blocks and phosphine-functionalized coupling partners employed in the miniaturized traceless Staudinger ligation platform for the synthesis of BRD4-targeting PROTACs. The used E3 ligands adducts target the E3 ligases cereblon (CRBN, upper panel) and Van Hippel Lindau (VHL, lower panel).

Solvent screening revealed substantial variation in conversion depending on both solvent composition and the azide substrate (Figure 2B, 2F). DMSO-containing solvent mixtures provided useful reaction media. For selected model reactions (**73** with azide **18**), conversions of approximately 70–73% were achieved in several mixed-solvent systems, whereas other substrate combinations showed substantially lower conversion under comparable conditions. The pronounced substrate dependence of the ligation highlights the importance of accounting for variable reaction conversions when interpreting D2B screening results.

**Figure 2.**
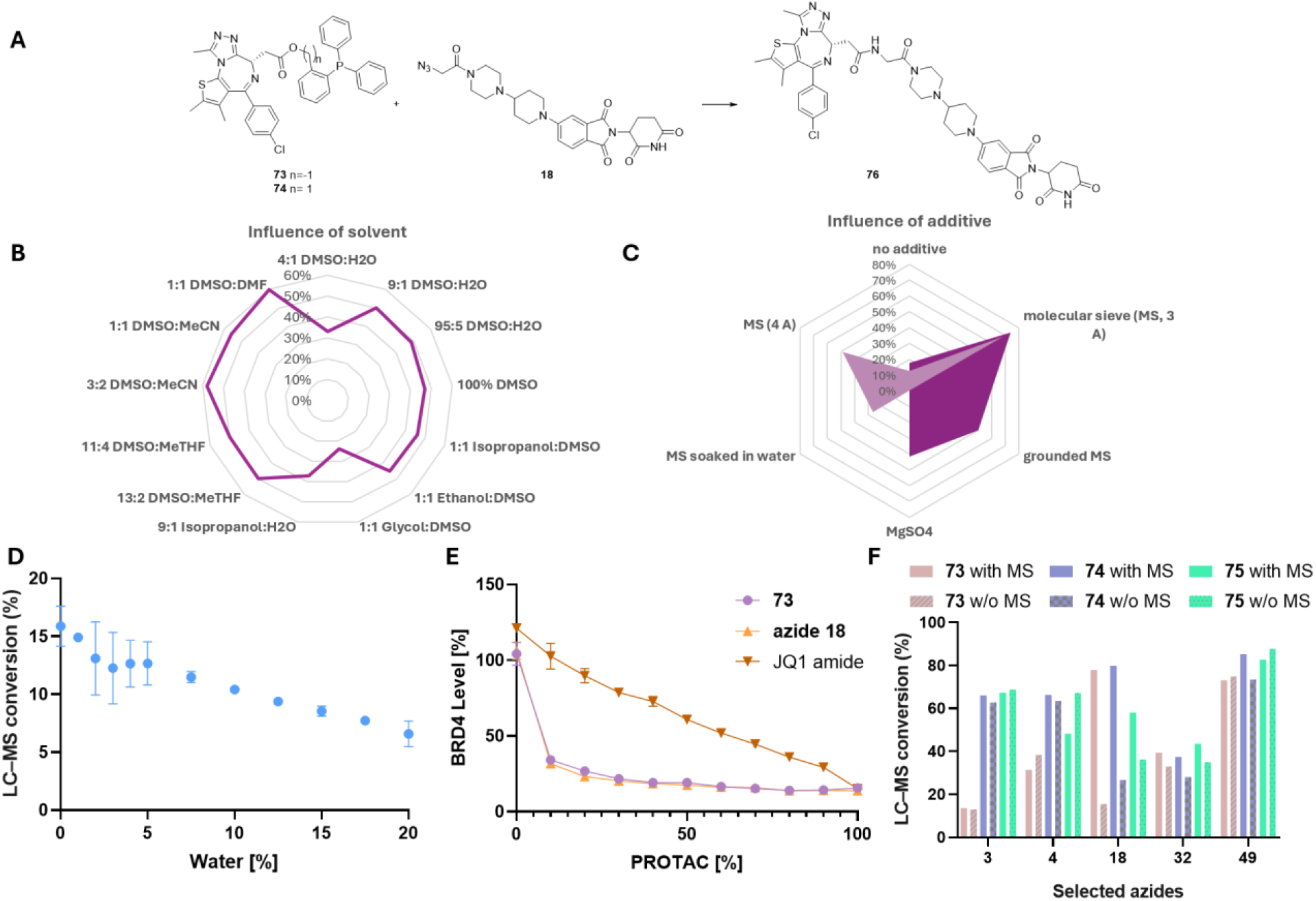
Optimization and evaluation of miniaturized traceless Staudinger ligation. (A) Reaction scheme. (B) Effect of solvent composition on LC–MS conversion (5 µL, 18 h, 40 °C). (C) Effect of dehydrating additives on LC–MS conversion (light purple: phosphine **73** in DMSO, 5 µL, 18 h; dark purple: DMSO/MeCN 3:2, 5 µL, 33 h). (D) Effect of added water on LC–MS conversion of phosphine **73** with azide **18** (DMF/DMSO 1:1, 5 µL). (E) Spiking experiments with PROTAC **76**, starting materials, and the JQ1-derived amide side product. (F) LC–MS conversion of five representative azides with phosphines **73–75** in the presence (solid bars) or absence (patterned bars) of molecular sieves (DMF/DMSO 1:1, 5 µL).

Because hydrolysis is required to complete the traceless Staudinger ligation, we investigated the influence of water on reaction conversion. Interestingly, we observed that azide stock solutions in DMSO that had undergone several freeze–thaw cycles generally afforded higher conversions than freshly prepared solutions of the same azides. We hypothesized that the repeated handling of these stock solutions might have led to the accumulation of trace amounts of water, which could facilitate the hydrolysis step. However, a systematic investigation of the water content revealed that increasing the proportion of added water reduced product conversion under the conditions examined (Figure 2D). We therefore explored whether separating the initial ligation from the subsequent hydrolysis step could improve yields of product formation. Following an initial reaction time of 17 h, water-containing mixtures were added, and the reactions were incubated for an additional 6 h. This approach, however, did not appreciably improve conversion (Table S3). Although base-promoted hydrolysis has been reported for related Staudinger ligation systems,^10^ we deliberately avoided the addition of a base to minimize the introduction of further components that could interfere with subsequent cellular screening of the crude reaction mixtures.

Given the apparent sensitivity of the reaction to water content, we next investigated whether drying agents could provide more consistent reaction conditions. Molecular sieves and MgSO_4_ were evaluated using a representative phosphine–azide combination. Intact molecular sieves markedly increased LC–MS conversion from 19% to 74%, whereas ground molecular sieves and MgSO_4_ afforded smaller improvements, reaching 48% and 43% conversion, respectively (Figure 2C). Interestingly, the superior performance of intact molecular sieves suggested that their influence may extend beyond the removal of residual water. Differences in surface properties, adsorption, or interactions between reaction intermediates and the solid additive could contribute to the observed effects^14^, although the underlying mechanism remains unresolved. To assess the generality of this effect, five representative azides were selected, comprising three α-azido ketones (**18, 49**, and **32**), an alkyl azide (**3**), and a PEG-containing azide (**4**). These were reacted with three phosphine-functionalized JQ1 derivatives (**73, 74**, and **75**) in DMF/DMSO (1:1), with and without molecular sieves (Figure 2F). Pronounced substrate-dependent differences were observed, even among azides belonging to the same structural class. For azide **18**, molecular sieves increased LC–MS conversion from 16% to 78% with **73**, from 31% to 80% with **74**, and from 36% to 58% with **75**. In contrast, azide **49** generally afforded high conversions irrespective of the presence of molecular sieves. These findings demonstrate that molecular sieve additives do not universally improve reaction outcomes, highlighting that their efficacy cannot be reliably predicted from the azide functional class. However, as its presence was advantageous in most cases, molecular sieves were retained in the screening workflow. Rather, ligation efficiency depends on the specific combination of azide, phosphine-functionalized scaffold, and reaction conditions. This substrate dependence highlights an important consideration for D2B library synthesis, where conditions optimized for a single model reaction may not translate directly to a structurally diverse collection of building blocks. Collectively, these experiments identified the principal challenges of transferring TSL to a miniaturized D2B format: solvent-dependent reagent solubility, accurate low-volume dispensing, and substrate-dependent conversion. Rather than pursuing extensive optimization for individual azides, we sought conditions that provided a practical balance between reaction performance, operational simplicity, and compatibility with direct biological evaluation.

A central requirement for D2B screening is that residual reagents, solvents, and reaction by-products do not substantially distort the biological readout. We therefore investigated the compatibility of representative reaction components with the cellular BRD4 degradation assay before interpreting the screening results. DMF, evaluated at the concentration present in the biological assay, showed no detectable cytotoxicity or BRD4 degradation in the HiBiT lytic assay. The phosphine-containing starting material and a representative azide were also examined in cytotoxicity and degradation experiments. Spiking experiments with PROTAC **76** showed that these starting materials did not appreciably alter PROTAC-induced BRD4 degradation under the investigated conditions (Figure 2E). In contrast, a JQ1-derived amide side product, detected exclusively in reactions involving the phenyl ester-containing phosphine **73**, attenuated PROTAC-induced BRD4 degradation in spiking experiments (Figure 2B). This effect is likely due to competition for BRD4 binding. Notably, the side product was not detected with the benzylic ester-containing phosphine **74**, highlighting the influence of phosphine ester structure on reaction chemoselectivity.^9^

Following reaction optimization and assessment of assay compatibility, the miniaturized TSL workflow was extended to the synthesis and screening of a broader library of BRD4-targeting PROTAC candidates. For this purpose, 72 azide building blocks were selected from our expanded in-house azide collection (Figure 1) and combined with phosphine-functionalized JQ1 derivatives. This strategy enabled exploration of amide-linked PROTAC architecture while retaining access to azide building blocks previously investigated in CuAAC-based D2B synthesis. The library synthesis afforded a mean LC–MS conversion of approximately 65%. The crude reaction mixtures were subsequently diluted and evaluated directly in a cellular BRD4 lytic HiBiT assay. Applying a threshold of ≤40% remaining BRD4, 23 screening hits were identified, with 19 originating from phosphine **74** and four from phosphine **75** (Figure 3A). To validate the D2B screening results, two selected PROTACs were synthesized, purified (Figure 3B), and characterized in concentration-dependent cellular BRD4 degradation assays. Both compounds were derived from the same azide building block (azide **18**) but differed in the phosphine-functionalized JQ1 precursor used for their synthesis. Following 24 h of cellular incubation, **76**, derived from phosphine **74**, exhibited a pDC_50_ of 8.2 and a D_max_ of 90%, whereas **77**, synthesized from the longer-linker precursor **75**, displayed a pDC_50_ of 7.0 and a D_max_ of 96%. Rescue experiments with the proteasome inhibitor Bortezomib and neddylation inhibitor MLN4924 restored BRD4 levels, as expected.

**Figure 3.**
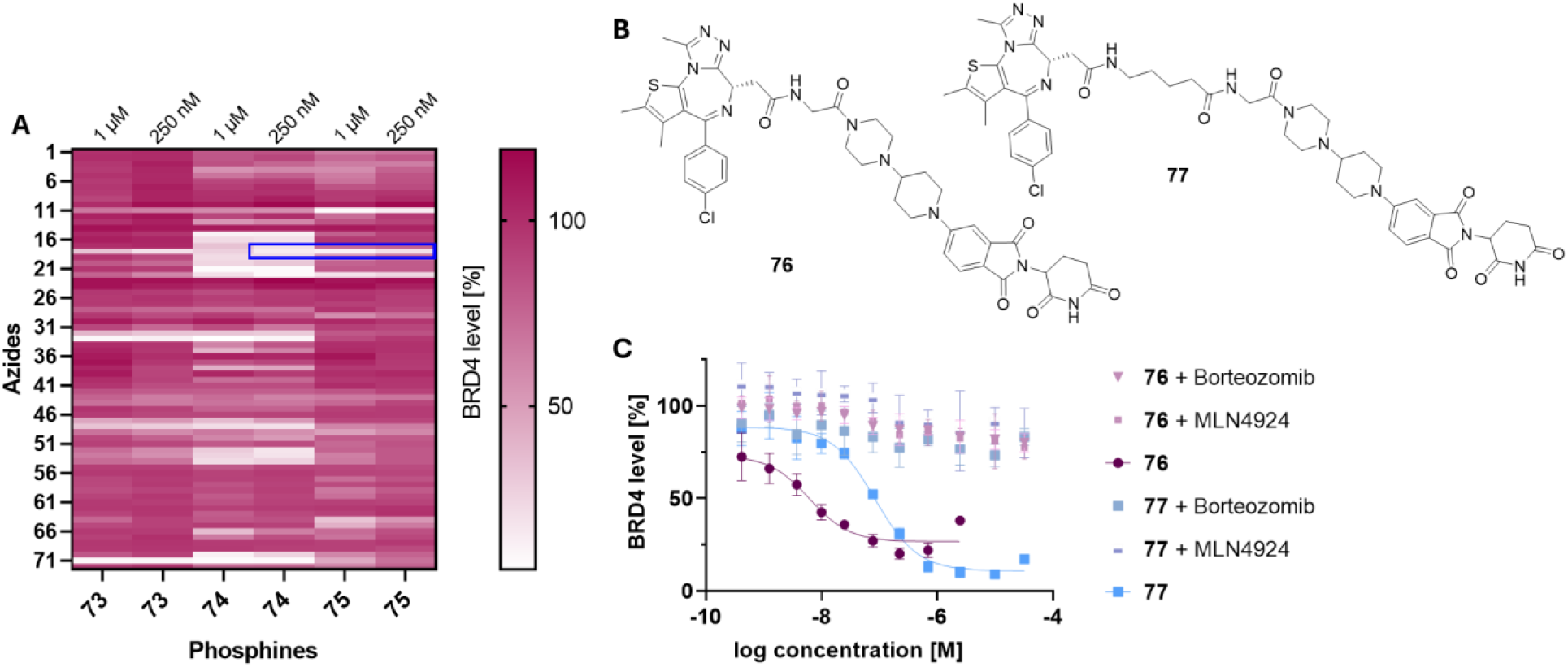
(A) BRD4 degradation screening of crude TSL reaction mixtures at different PROTAC concentrations using the lytic HiBiT assay (6 h incubation); the blue box highlights selected hits. (B) Chemical structures of the selected PROTACs **76** and **77.** (C) Dose–response and rescue experiments with purified PROTACs **76** and **77** following 6 h of cellular incubation.

Overall, this work establishes miniaturized, base-free traceless Staudinger ligation as a reliable D2B strategy for the discovery of amide-linked BRD4-targeting PROTACs, yielding multiple screening hits and a highly potent validated degrader (**76**). Complementing our previously established CuAAC-based workflow, TSL enables the exploration of distinct linker architectures using a shared azide collection, thereby broadening the accessible chemical space for PROTAC discovery. Notably, the shorter-linker series derived from phosphine **74** yielded substantially more screening hits than the longer-linker series derived from **75** (19 versus 4), suggesting that shorter linkers may be advantageous for BRD4 degradation within the investigated library.

## Acknowledgements

The authors acknowledge funding by the Structural Genomics Consortium (SGC), a registered charity (no: 1097737) that receives direct member funding from Amgen Inc., Janssen Pharmaceutica NV, and Bristol-Myers Squibb Company, as well as grant funding from the Innovative Health Initiative Joint Undertaking (IHI JU; LIGAND-AI, grant agreement No. 101252959) and by the German Cancer Aid (DKH, TACTIC-70115201). The figures were created with BioRender.com.

